# Hippocampal sequences traverse a memory space

**DOI:** 10.1101/2025.11.21.689701

**Authors:** Olivier C. Ulrich, Daniel Molinuevo-Gomez, Margaret R. Lane, James B. Priestley

## Abstract

Highly reliable sequential dynamics organize neural activity in the hippocampus and progress with changes in physical, sensory, or cognitive variables, typified by place cells during active exploration. But sensory and behavioral variables are often confounded, and so it remains intensely debated whether hippocampal sequences are driven by sensory content directly or rather reflect the alignment of internally generated dynamics to external perception. Here we used mouse virtual reality to characterize hippocampal dynamics in environments of systematically different sensory complexity, and to test the robustness of neural sequences to manipulations of exact sensory content. The structure of neural sequences was perturbed by changes in both moment-to-moment sensation and ongoing behavior, in a pattern that was best explained by the different memory strategies employed by the animal in each context. Neural subsequences could be flexibly interrupted, inserted, and resumed wherever a memory of prior experience was violated, indicating that the propagation of hippocampal dynamics was not rigidly fixed by internal networks and initial conditions. Rather, we argue that hippocampal sequences explore an abstract memory space of selectively remembered experiences, which flexibly encompasses both sensory and internal factors in a context-dependent manner.

## Introduction

It is now widely understood that hippocampal representations reflect more than strictly a spatial map of the environment [1, 2]. The circuit responds to a diverse set of sensory variables decoupled from physical position, including odors [3, 4], tones [5], visual features [6–8], object identity [9, 10], and more cognitive variables such as reward [11, 12]. Because these variables can often be arranged as a continuous stimulus space, it has been suggested that this polymodal coding reflects a more general “mapping” function on sensory and cognitive spaces [13, 14]. Compatible with this idea, many models explain place coding through compression of multi-sensory information during experience, where place fields generically emerge as a solution to balancing representation sparsity [15, 16] or neural independence [17–19] with the underlying smoothness and redundancy of sensation during movement. This is consistent with experimental observations of heterogeneous density and precision of hippocampal place codes, which can co-vary with the density of sensory cues [20] and the rate of perceptual changes during movement [21, 22].

But there is considerable debate as to whether these responses reflect bona fide coding for external variables (“place cells”, “tone cells”), or rather an artifactual alignment of sensory experience to internally generated neural activity [23–25]. Internally generated neural sequences are hypothesized to arise through autonomous network dynamics in the hippocampus [26], initiated or driven by other cognitive signals such as action plans [27] that are partially confounded with cues in the animal’s physical environment during stereotyped behavior. Accordingly, numerous studies have reported the sensitivity of hippocampal place fields to reward and behavioral goals [11, 12, 27–29], and the presence of place cell sequences can depend strongly on continuous task engagement [30]. But hippocampal representations are known to dynamically adapt to changing environmental factors, such as manipulation of salient sensory cues [3, 8, 31, 32], suggesting that these neural sequences cannot be strictly constrained by preconfigured network structure and internal dynamics alone.

Although there have been many conceptual and formal attempts to reconcile these accounts under a unified theory of hippocampal function [13, 15, 33–39], it remains unresolved precisely what place cells encode [26]. Here we sought to assess the control of hippocampal sequences by sensory and internal factors, through the design of virtual reality (VR) tasks that allowed us to control the complexity of the animals’ sensory environment, and to systematically perturb patterns of sensory information during goal-directed behavior. To this end, we built minimally structured VR environments using samples of procedurally generated textures [40, 41]. These parametrically controllable environments allowed us to ask both how gross statistical features of the visual sensory environment affected the neural representation of space, and also how ongoing neural sequences react to visual perturbations. Combining VR paradigms with large scale optical neural recordings [42, 43], we tested the robustness of ongoing hippocampal dynamics to changes in external sensory variables and behavior. By replacing or inserting unique but statistically similar sensory content in learned environments while spatial goals remained unchanged, we report differential alignment of neural activity to sensation and action that reflected animals’ divergent memory strategies across sensory contexts.

## Results

### Behavioral and neural discrimination in procedurally generated virtual environments

We developed a visual VR system (Fig. 1A) and an automated procedure for generating virtual linear tracks with statistically homogeneous visual features [40–42], namely characterized by a consistent visual scale of the textures on the walls. Using this system, we taught mice a simple contextual discrimination task using 2 environments of equal length (4 m) but created with different visual statistics (wall textures containing high or low frequency frozen noise patterns; the pattern for each context does not change across trials, Fig. 1A, B, S1). On each trial, mice obtained sucrose water rewards by licking at the correct goal zone for the current context (Fig. 1B; Near = 2.5 m or Far = 3.4 m; goal locations were randomly assigned to High and Low contexts across mice). Once trained, mice reliably restricted their licking to locations in advance of the correct goal location (Fig. 1C,F) until reaching satiety (Fig. S1). Similarly, the mice developed context-specific preparatory slowing, decreasing their velocity in advance of the correct goal location (Fig. 1D,E). Together, licking behavior and velocity before the goal zones could be used to accurately decode the identity of the current VR environment at both potential goal locations (Fig. 1G), indicating that the mice used at least the global context statistics (high or low frequency visual textures) to guide behavior on each trial.

**Figure 1.**
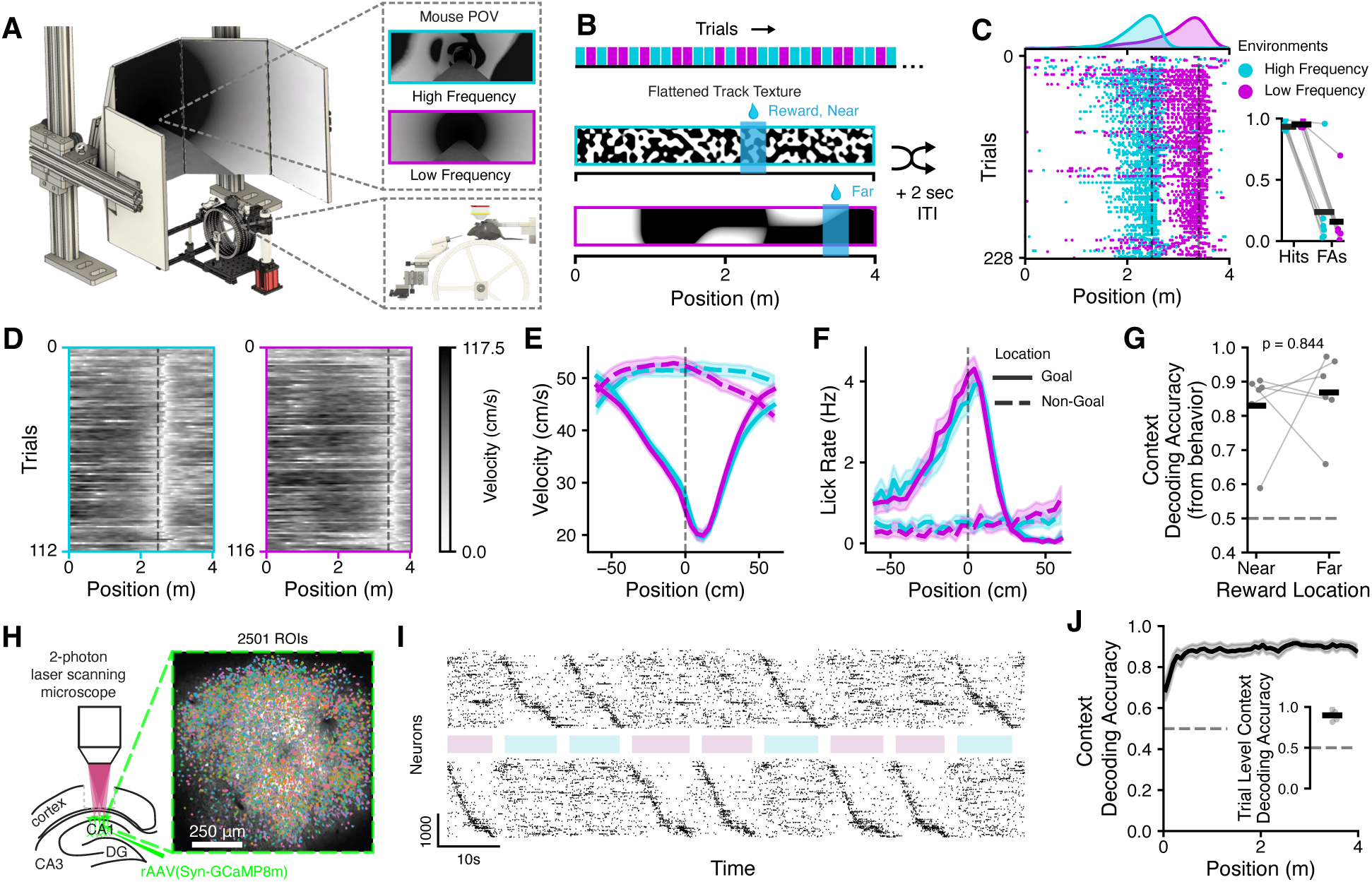
Behavioral and neural discrimination in minimally structured virtual reality environments. **A:** Head-fixed virtual reality (VR) system for mice, and example procedurally generated VR environments. **B:** Trials alternated pseudorandomly between two environments with different visual statistics. The mapping of goal zone locations (Near/Far) to context (High/Low) were randomized across mice. **C:** Raster plot of licking data in the context discrimination task. Inset: hit rate (fraction rewards collected) and false alarm rate (fraction of trials with a lick in the incorrect goal zone) in each environment. **D:** Mouse velocity for the example session shown in **C**, plotted separately by context. **E:** Average velocity at each distance relative to the goal (or non-goal) zones (non-goal is the position of the goal in the other context). **F:** As in E, for lick rate, with context-specific preparatory licking. **G:** Context decoding accuracy obtained via logistic regression from behavior data (velocity and lick rate) 28 cm before the early or late goal zones (n = 6 mice, Wilcoxon signed rank test). **H:** Schematic of 2-photon imaging preparation (left), and example field-of-view (right). **I:** Population activity across trials during the task. Unique sequences of place cells tile each environment. Neurons are sorted by the latency of peak spatial activity in the high (top) or low (bottom) frequency context. **J:** Accuracy of linear SVM decoder trained on the neural data to predict context at each distance in VR. Inset: trial-level accuracy obtained by pooling classifier votes across positions on each trial. Error bars are mean *±* s.e.m.

Mice were stereotaxically injected with an rAAV vector to drive expression of the fluorescence calcium indicator *GCaMP8m* in CA1 sub-region of the dorsal hippocampus. Optical access to the hippocampus was facilitated by implanting a chronic imaging window immediately above the external capsule (Fig. 1H). Using a custom 2-photon microscope, we recorded up to 3095 CA1 neurons simultaneously during VR behavior (1297 ± 216 ROIs per session, mean ± s.e.m., Fig. S2). Spatial masks and neuropil-subtracted fluorescence traces were extracted using Suite2p [44], and the resulting traces were deconvolved to reduce the impact of the intrinsic autocorrelation of the calcium indicator [45]. Unique neural sequences were readily observed in the sorted population activity in each context (Fig. 1I) with complete coverage of the virtual environments. As a result, we could accurately decode the identity of the current VR context (High or Low) at every matched distance within the two environments from the neural activity (Fig. 1J).

### Place coding resolution does not follow the spatial scale of visual features

We first asked whether there were gross differences in the structure of the neural code in each context that reflected the different visual statistics of the virtual environments. Place fields tiled the track in both contexts (Fig. 1I, 2A). Following previous work on sensory richness [20, 21], we expected that spatial coding would appear more precise or dense in the High environment, but were surprised to find that spatial tuning of single neurons was generally more precise and reliable in the Low environment (Fig. 2C). The population spatial autocorrelation was also slightly wider in the High context (Fig. 2D), indicative of lower spatial resolution. We further quantified the precision of spatial coding in each environment by training decoders to discriminate between each pair of positions throughout the track (Fig. 2E), and while performance was generally highly accurate across all pairs, the accuracy in the High context was slightly worse across most distances (Fig. 2F). We observed a similar trend in most mice when using the ensemble of pairwise classifiers to decode position (Fig. 2G). In sum, we observe robust spatial coding in both contexts, but the neural code in the Low context is slightly more precise, despite the wider spatial scale of the sensory stimuli in the environment. These data indicate that, at least in these relatively abstract, minimally structured environments, the spatial autocorrelation of the neural representation is not directly inherited from the correlation structure of the sensory stimuli.

**Figure 2.**
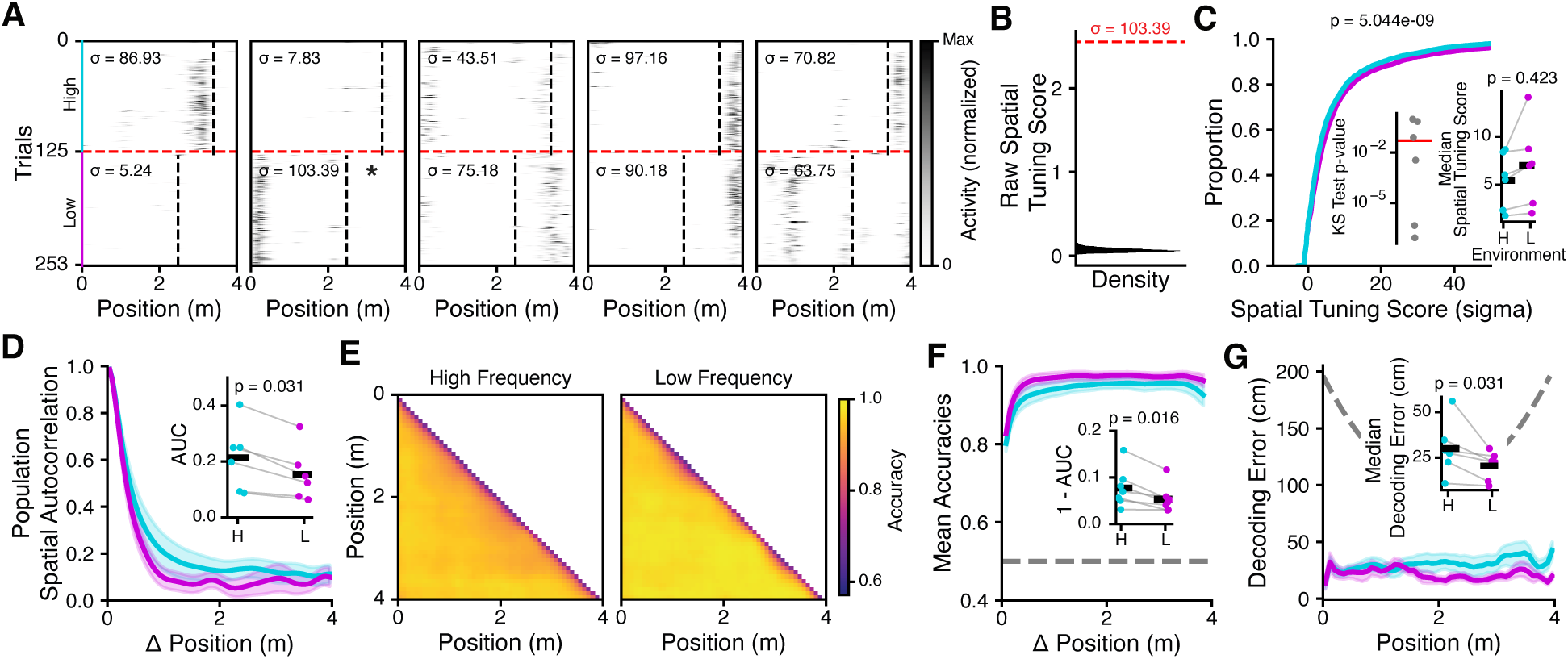
Resolution of position coding is paradoxically lower in environments of finer visuospatial scale. **A:** Example single neuron spatial tuning across trials in each environment (spatial tuning score inset). **B:** Spatial tuning score and null distribution computed for the 2nd example in **A**. The null distribution is generated by independently shifting the spatial tuning profile on each trial by a random distance before recalculating the trial-averaged tuning curve and spatial tuning score. The normalized tuning score *σ* for each neuron is standardized relative to the null distribution. **C:** Cumulative distribution of spatial tuning scores in each environment (n = 12597 neurons, KS test). Inset left: KS test p-value for each mouse comparing High vs Low scores. Inset right: median tuning scores across each environment (n = 6 mice, Wilcoxon signed rank test). **D:** Population vector spatial autocorrelation in each environment. Inset: area under the curve (AUC, Wilcoxon signed rank test). **E:** Matrix of accuracy scores of pairwise decoders (linear SVM) trained to discriminate each pair of positions in the environment, shown for each context. **F:** Summary of pairwise classifier performance in **E** at each distance in the environment. Inset: 1 - AUC (Wilcoxon signed rank test). **G:** Mean absolute decoding error at each position in the environment. Inset: average error across positions. All error bars are mean *±* s.e.m.

### Neural sequences relative to goals are largely orthogonalized by context

Because we did not find a straightforward connection to the sensory parameters of the environment, we wondered whether other aspects of the task affected the observed neural representations. Reward- and goal-related signals are pervasive in many hippocampal recordings and across diverse behavioral paradigms [27], with some reports of context-invariant goal coding [11] even at substantial distances relative to goal locations [12, 29]. We took the perspective of a downstream reader and asked how well we could decode the relative distance to goals from hippocampal population activity. In particular, we studied the cross-condition generalization performance (CCGP) of decoders trained to predict distance-to-goal in one environment, by testing the classifiers’ performance in the other environment. This analysis provides geometric insight about the neural representation format and its capacity for abstraction and generalization across contexts [46]. We used this analysis to distinguish representational formats such as converging sequences to common goal cells, parallel sequences that abstractly encode distance-to-goal, and independent sequences that encode the goal differently in each context (Fig. 3A).

**Figure 3.**
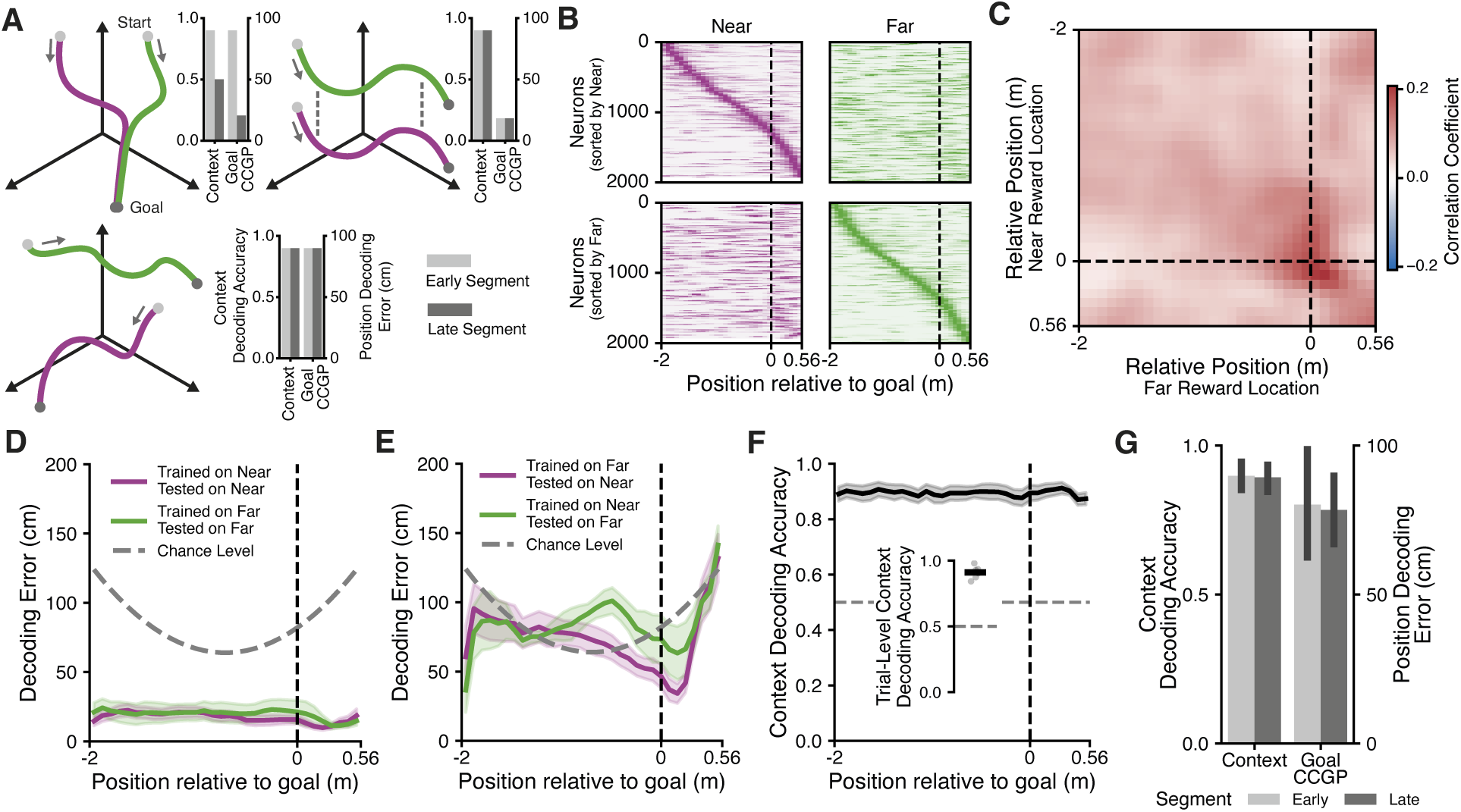
Limited evidence for context-invariant goal coding. **A:** State space representations of 3 hypothetical coding scenarios. Upper left: independent place cell sequences in each context converge to a common, inseparable goal-coding population as the animal approaches the goal zones. Upper right: place cell sequences are separable throughout the environment but aligned (“abstract” representation of distance-to-goal). Bottom left: place cell sequences in each context are orthogonal. Each scenario makes different predictions regarding the decodability of context and the generalization of distance-to-goal decoders across contexts (CCGP). **B:** Example population neural sequences for trials where the Near (left) or Far (right) goal location was rewarded, shown sorted by the location of peak activity in each trial type. **C**: Population vector cross-correlation of neural spatial tuning curves, in goal-relative space, in Near-rewarded vs Far-rewarded trials. There is a small increase in correlation around the location of the goal zones between contexts. **D:** Spatial decoding error as in 2G (linear SVM), in goal-relative space. **E:** Cross-condition generalization performance of spatial position decoders trained on Near trials and tested on Far trials (green) or the reverse (purple). Decoders fail to generalize across contexts except in a very spatially limited region close to the goal zone. **F:** Context decoding accuracy at each position in goal-relative space. Context decoding remains highly accurate close to the goal zones, indicating the population representation of the goal is highly separable between contexts. **G:** Summary of decoding results. Context decoding is highly accurate, while goal-relative position decoding CCGP exhibits high errors, indicating mainly independent representations. n = 6 mice, all error bars are mean *±* s.e.m.

Comparing the alignment of neural sequences during trials where the goal was located in the Near or Far position (Fig. 3B), we did not see an obvious structure that generalized across trial types. Analyzing the population vector cross-correlations between Near and Far trials, we observed a small increased correlation between the location of the goal zone in each context (Fig. 3C), but this effect was spatially restricted to the goal location itself. As in Fig. 2, we trained spatial decoders to predict the position of the animal, but now in goal-relative coordinates, and tested the decoders both within (Fig. 3D) and across (CCGP, Fig. 3E) trial types. Decoders largely failed to generalize across trial types, performing at chance level except at positions very close to the goal zone (Fig. 3E), consistent with the increased correlation around the goal zone (Fig. 3C). Simultaneously, we could accurately decode the trial type independently at all goal-relative positions (Fig. 3F, 3G). Combined, these results indicate that the neural representations in the two contexts were mainly independent.

### Conditional integration of altered sensory stimuli

The previous analysis indicated that neural representations in our VR task were strongly context dependent, but the structure of those representations was not obviously related to the parameters of the visual environment itself (beyond encoding the global context identity, High or Low). One possibility is that the neural sequence still does not encode the identity of the sensory content within the environments, but rather arises from the animals’ internal action plan [27], which in this case is context dependent (i.e., the distance-to-goal differs between environments). We next tested whether the observed sequences were responsive to manipulations of the sensory content on the environment walls. We performed the same context discrimination as before (Fig. 1), but now in 50% of trials in each context, we replaced the visual texture on the walls in the 2nd half of the environment (including the locations of the goal zones). The manipulation is not immediately visible to the animal from the start of the track, due to in-game lighting effects that limit the animals’ line-of-sight to 1 m ahead of their current position in the VR. The altered visual stimulus was generated using the same procedural noise algorithm [41] and was matched to the visual parameters of the original environment, so each track maintained roughly the same image statistics. The same altered stimulus was shown on each Replaced trial in each context.

For a strictly distance-based memory [25, 27], the altered sensory cues are irrelevant to the task; the sequence at the beginning of the track already disambiguated the context, and the altered sensory cues are not yet visible. An internally generated sequence could then be primed by the initial conditions that determine the trial type (context), and progress autonomously to the correct goal distance [27]. Nevertheless, we observed the appearance of many new place fields when animals explored the replaced section of the track (Fig. 4B). We inspected the population vector spatial correlations between the Original and Replaced versions of the track, and found a strong neural decorrelation in the last 2 meters of the environment, where the sensory information had changed, but only in the Low context (Fig. 4C-E). Strikingly, the sequence in the High environment remained largely unchanged, except in the positions at the very end of the environment. In both cases, the quality of spatial coding was similar between the Original and Replaced trials in the perturbed region of the environments (Fig. S4). Consistent with the spatially restricted neural decorrelation, we could decode the Original vs Replaced visual environment but only in the 2nd half of the environment and only in the Low context (Fig. 4F,G).

**Figure 4.**
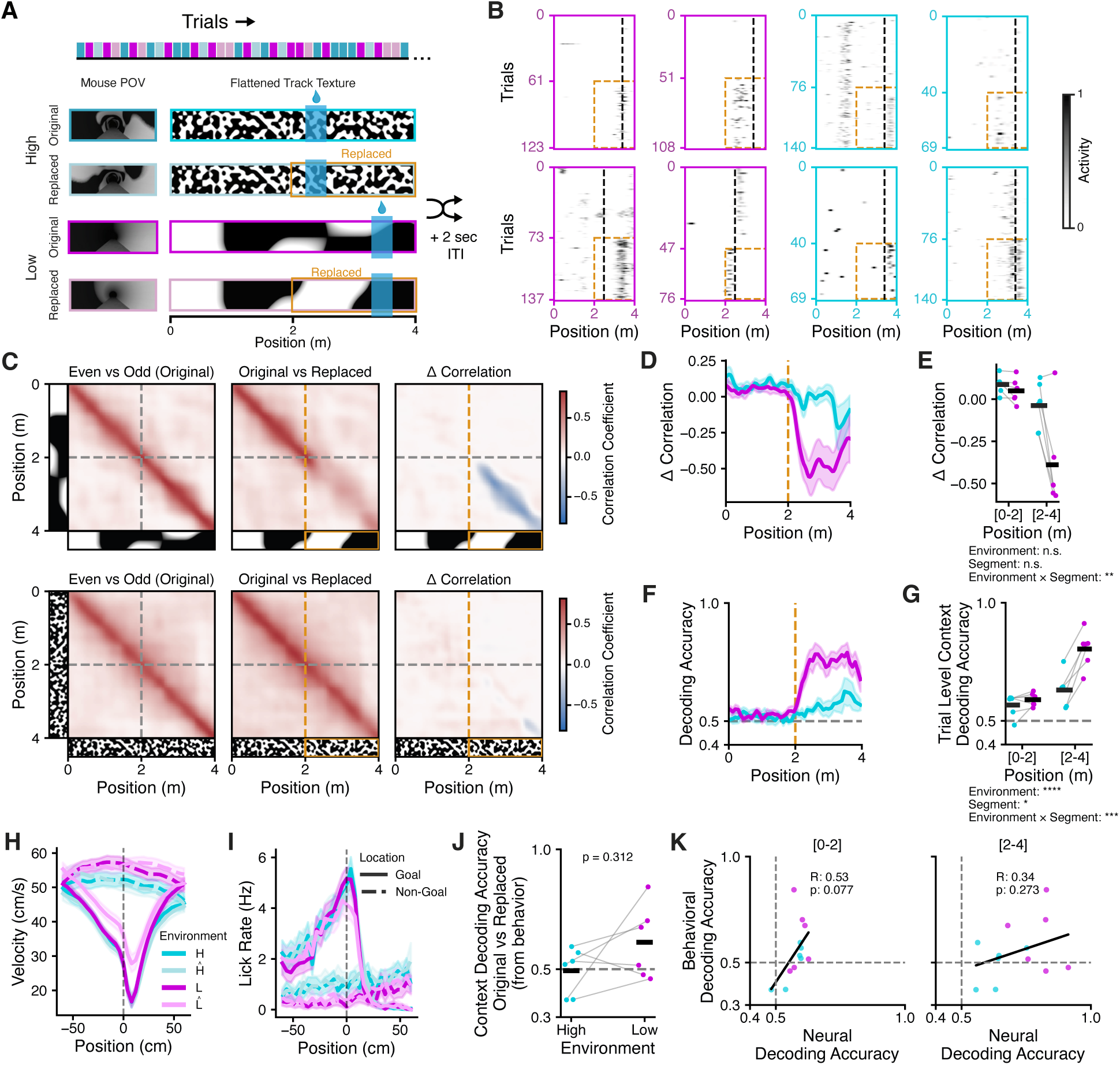
Neural sequences diverge when sensory content is altered, but only in low complexity environments. **A:** Schematic of task design. Mice performed the same context discrimination paradigm as before, but on 50% of trials, the visual texture on the walls was replaced (yellow highlighted region) with a different pattern with identical statistics (i.e., high or low frequency Perlin noise). The goal zone in Replaced trials remained in the same location (i.e. distance relative to the start of the trial). **B:** Example single neuron spatial tuning within a single environment, sorted by Original vs Replaced trials (the replaced section of the environment is highlighted in yellow). The replaced texture evokes different patterns of place field firing. Note the absolute distance to goal (black dashed line) does not change. **C:** Population vector cross-correlations for each pair of spatial positions (averaged across mice), between Even vs Odd trials in the Original environment (left), between Original vs Replaced trials (center), and their difference (right), shown for the Low (top) and High (bottom) contexts. The replacement texture reliably decorrelates the 2nd half of the Low context. **D:** Summary of the diagonal of the Δ correlation matrices plotted in **C** (right) for each mouse. **E:** Average Δ correlation in **D** for each mouse and context, shown for the first half (0-2 m) and second half (2-4 m) of the environment (linear mixed effects model). **F:** Decoding accuracy (linear SVM) from classifiers trained to predict Original vs Replaced trials in each context. **G:** As in **E**, but summarizing the decoding results from **F** (linear mixed effects model). **H:** Velocity around the goal zone and non-goal zones for all 4 trial types (Original = *H, L*; Replaced = *H^^^, L^^^*). **I:** As in **H**, but for lick rate. **J:** Accuracy of logistic regression trained to predict Original vs Replaced trials from behavior data (pre-goal velocity and licking, Wilcoxon signed rank test). **K:** Correlation between neural decoding accuracy in **F** and behavior decoding accuracy in **J**, shown separately for 0-2 m and 2-4 m. Error bars are mean *±* s.e.m.

Overall, these data indicated that there was a context-dependent change in the structure of the neural sequence: manipulating sensory cues in the Low environment drove a bifurcation in the neural representation that encoded the altered sensory experience, whereas the neural code in the High environment remained unchanged across conditions. We asked whether this dichotomy was also evident in the velocity and licking behavior of the animals across each context and trial type (Fig. 4H, I). We observed slightly increased running speed in the Low context during replaced trials, but overall the behavior differences between contexts were generally low and our ability to decode Original from Replaced trials from behavior near the Goal zones was poor (Fig. 4J). Behavioral decoding accuracy was also not significantly correlated with neural decoding accuracy of Original vs Replaced trials (Fig. 4K). Given that the neural decorrelation was caused by the appearance of new, localized place fields in the Replaced Low trials (Fig. 4B-G), the neural decoding cannot be explained as a simple function of the animals’ small change in average velocity (Fig. 4H). Notably the distance-to-goal remained unchanged across the sensory manipulations in both contexts, and so it is difficult to disambiguate the precise roles of behavior and sensation in the observed context-dependent remapping [4, 47].

### Neural sequences are aligned to context-specific memory content

To resolve the factors that controlled remapping across contexts, we devised a second sensory manipulation where the sensory cues around the goal locations now remained fixed. Instead, in 50% of trials in each context, we inserted an additional 2 m track segment with matching visual statistics [41] as before (Fig. 5A), such that in half of all trials, the environment was now 6 m in length and the distance to the goal locations in these Inserted trials was increased by 2 m. Once again, we found that a new sequence of place fields emerged in the Inserted track segment (Fig. 5B). Neurons whose place fields were located between 2-4 m in the Original trials also shifted their activity by 2 m in the Inserted trials, such that they were aligned to the same visual stimuli (Fig. 5B). This pattern of remapping was easily visible in the population vector cross-correlation between Original and Inserted trials (Fig. 5D-F), where the Original sequence was interrupted specifically between 2-4 m in Inserted trials, where the additional track segment was located. We similarly observed that the quality of spatial coding was similar between Original and Inserted trials (Fig. S4), indicating the observed decorrelation was not driven by a reduction in spatially modulated activity. The neural activity was then realigned between Original trials from 2-4 m and Inserted trials from 4-6 m (i.e., where the visual input was matched). Notably, and unlike the replacement experiments in Fig. 4, here we found a decorrelation in the manipulated track segment in both the Low and High contexts. As in Fig. 4, the representation quickly decorrelated in the Low context immediately at 2 m, where the Inserted track segment and altered sensory content began (Fig. 5D-F). But the decorrelation in the High context appeared delayed, with the original sequence maintained for a small distance within the Inserted track segment (Fig. 5D,E). While in this experiment the representation in the High context also diverged in the altered track segment, it appeared that the decorrelation was not strictly aligned to the transition to different sensory content.

**Figure 5.**
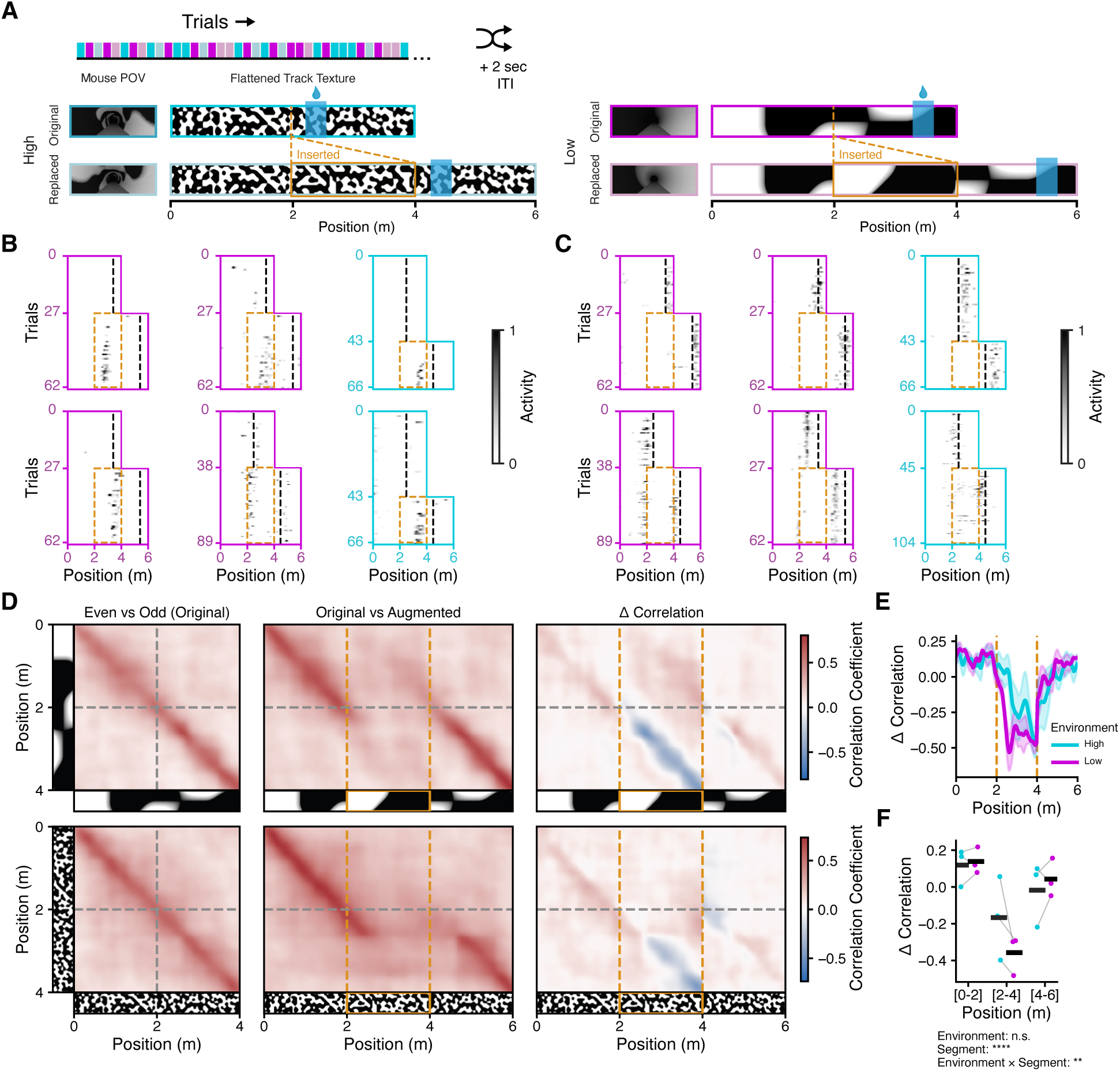
Ongoing neural sequences can be interrupted and resumed by inserting new track segments into the environment. **A:** Schematic of task design. Mice performed the same context discrimination paradigm as before, but on 50% of trials in each context, the virtual environment was augmented by inserting an additional 2 m segment (yellow highlighted region) which contained a different visual pattern with identical statistics (i.e., high or low frequency Perlin noise). Goal locations remained in the same relative location within the original second half of the environment, now located at 4-6 m in the Inserted trials. relative to the start of the trial). **B:** Example single neuron spatial tuning within a single environment, sorted by Original vs Inserted trials (the inserted section of the environment is highlighted in yellow). The inserted track segment is populated with its own sequence of place cells. Note the absolute distance to goal (black dashed line) does not change relative to the last 2 m of the environment on each trial, but is shifted 2 m back relative to the start of the trial in the Inserted trials. **C:** As in **B**, but for example neurons whose place field was originally located between 2-4 m and shifted their response by 2 m in the inserted trials. **D:** Population vector cross-correlations for each pair of spatial positions computed as in Fig. 4C (averaged across mice). In the Δ correlation plots, the difference for positions 4-6 m is computed relative to 2-4 m in the left plot. The original sequence is reliably interrupted by and resumed after the inserted track segment. **E:** Summary of the diagonal of the Δ correlation matrices plotted in **C** (right) for each environment. **F:** Average Δ correlation in **D** for each mouse and context, shown for the original first half (0-2 m), inserted segment (2-4 m), and original second half (4-6 m) of the environment (linear mixed effects model).

We further investigated the source of variability in the decorrelation pattern of the representation on Inserted trials. Specifically, we wondered if animals exhibited different behavior between the High and Low contexts on Original and Inserted trials (Fig. 6A, B). We compared animals’ preparatory slowing and licking in the region around the goal location in Original trials, and the “false” goal location at the equivalent distance during Inserted trials. While mice in the Low context generally avoided slowing and licking during the traversal of the inserted region of track, they exhibited erroneous behavior in the Inserted High trials, indicating they searched for expected reward at the previously memorized distance (Fig. 6A). We quantified this pattern by attempting to predict Original from Inserted trials from velocity and licking in the pre-goal and pre-false goal locations (Fig. 6C). This analysis was highly accurate during Low trials, indicating well separated behavior, but deteriorated in High trials, consistent with erroneous goal-seeking behavior. We repeated this analysis comparing behavior at the true goal locations in both Original (2-4 m) and Inserted (4 - 6 m) trials, where decoding was generally poor, indicating largely similar behavior (Fig. 6D). To summarize, it appeared that during each High Inserted trial, the mice would seek reward twice – first at the remembered distance (2-4 m), and then again in the final segment of the track after failing to obtain reward earlier.

**Figure 6.**
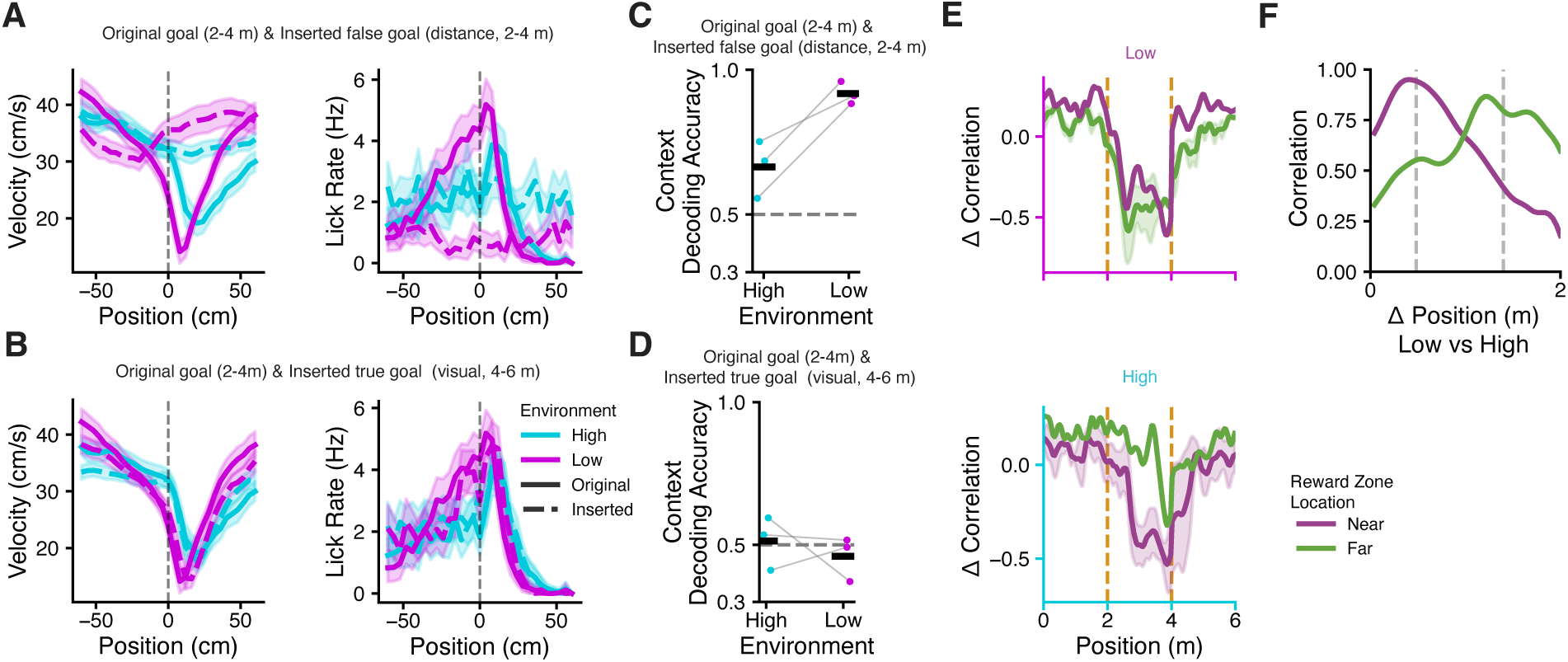
Sequence progression is differentially predicted by sensory content and behavior across contexts. **A:** Velocity and lick rate around the goal locations in Original trials and the “false” goal, located at the same distance in Inserted trials. In High trials, the mice exhibit similar preparatory slowing and licking for both Original and Inserted trials. **B:** As in **A**, but comparing the true goal locations in Original and Inserted trials (different distances). The behavior is largely similar. **C:** Decoding Original vs Inserted trials (logistic regression) from the pre-Goal velocity and licking behavior in **A**. Decoding accuracy is consistently worse in High trials, indicating the animals’ behavior is very similar at the Original goal location, and the equivalent distance in Inserted trials. **D:** As in **C**, but comparing behavior at the Original and Inserted true goal locations in **B**. Decoding accuracy is poor, indicating similar behavior profiles. **E:** As in Fig. 5E, but with data plotted separately for experiments where the High or Low context had the Near goal zone, or the Far goal zone. In the Low context (top), the neural representation quickly decorrelates in the inserted (2-4 m) segment of the track. In the High context, the decorrelation is delayed for Near trials and even more so for Far trials. **F:** Optimal alignment of the 2-4 m decorrelation pattern in **E** for High trials with different segments of Low trials. The gray lines indicate the expected peak alignment if decorrelation is shifted from the beginning of the Inserted region for Low trials to the two expected goal locations (+0.5 m and +1.4 m) for High trials. The data coincide with these expected intervals, indicating the High sequence is disrupted only after the animal fails to receive reward at the expected distance.

Does this behavior explain the delayed neural decorrelation (Fig. 5D-F) in High Inserted trials? We took advantage of the fact that the goal zone location was randomized between Near (2.5 m) or Far (3.4 m) locations across mice (Fig 1B). Plotting the decorrelation curves from Fig. 5E separately for Near and Far mice in each context, we found that the decorrelation occurred immediately at the boundary of the Inserted segment regardless of the Original goal location in the Low context. In other words, the neural transition was aligned to the change in sensory information. However, decorrelation was spatially delayed in the High context, in both Near and Far mice. The lag in decorrelation also corresponded specifically to the distance between the boundary of the

Inserted track segment (2 m) and the memorized goal distance (Δ = 0.5 m for Near, and 1.4 m for Far, 6F). Combined with the effects observed in Fig. 4, these experiments indicate that the mice remembered fundamentally different components of the experience in each context during the task. In the Low environment, the animals recalled the specific sensory stimuli in the environment, whereas in the High environment, they recalled the distance to the goal within the global context. This explains why altering the exact sensory content always evokes a detectable neural change in the Low environment, as it violates the animals’ prior memory of the experience (Fig. 4, 5, 6). In contrast, a neural change in the High environment is observed only when violating the animal’s remembered distance to the goal within the track.

## Discussion

There is growing consensus that the hippocampus does not truly comprise discrete functional cell types for different modalities of experience [26, 48]. Instead, a diversity of sensory and idiothetic responses exist within the same neurons and across the entire circuit in a widely distributed neural code [3, 48–52]. In line with our results, we argue that further progress toward a unified understanding of hippocampal function will be enabled by analyzing contradictory reports of what information may or may not be encoded by the circuit across diverse behaviors, contexts, and individuals. For example, we find very little evidence of abstract goal coding during our behavioral paradigms (Fig. 3) despite many earlier reports to the contrary [11, 12, 27], while we find context-dependent alignment of hippocampal sequences to moment-to-moment sensory content [7] or distance to goals (Fig. 4, 5, 6) [12, 27]. The resolution must lie in the different behavioral strategies and attentional biases of the animals across these paradigms [4, 30, 47], which are not always observable and likely confer different memory for these experiences. Our findings are congruent with earlier ideas that the hippocampal code reflects not a representation of strictly physical space, but rather an abstract memory space built through learning, which may selectively and variably include both task-relevant and irrelevant information [37].

Our main result demonstrates that the progression of hippocampal sequences differentially aligns to external or internal variables in environments of different visual complexity. This dichotomy is intimately linked to the different memory strategies employed by the animals for identifying the goal in each context. Sequences in the Low context are highly reactive to the injection of different sensory content, regardless of whether it coincides with the goal location (Fig. 4) or not (Fig. 5). In the Inserted environments, both the neural activity and animal behavior were yoked to specific visual experience, with the mice accurately identifying the goal zone at the shifted location without erroneous slowing or licking in the inserted region of the track (i.e. the original goal distances, Fig. 5, 6). Conversely, in the High context, we see no remapping when the sensory content is changed but the distance to the goal location remains the same (Fig. 4). When the track is augmented and the goal zone shifted in physical space, the original sequence continues uninterrupted until precisely the moment the animal fails to collect reward at the original, remembered distance (Fig. 5, 6).

While this activity pattern is consistent with a code for distance or progress to goal [12, 25, 27, 53], the High sequence also encodes the overall context identity (Fig. 1, 2) through a representation that is orthogonal to the Low environment (Fig. 1, 3). Sequence progression may align to an internal variable such as integrated self motion or time [23, 26, 54, 55], but these dynamics may arise via feedforward computation (e.g. on grid cell input) rather than local recurrent mechanisms in the hippocampus during active behavior [10, 36, 53, 55–58]. We interpret these data to mean that the animals generally do not remember the sensory details of specific locations in the High context, but they recognize the global statistics of the environment and memorize the goal location by distance. Path integration can be prone to accumulated errors [59], which could explain the paradoxically lower spatial precision of neural activity in the High environment compared to Low (Fig. 2). The sensory information in both contexts are safely within the perceptual resolution of the mouse visual system [60, 61], and mice can be trained to discriminate between two distinct High environments (Fig. S3). But in the High vs Low experiments, we speculate that the greater information content of the High context may bias animals to prefer distance estimation over visual template matching, to minimize working memory requirements. Future work can build on our parametric framework by systematically manipulating animal velocity (e.g. through VR gain modulation [22, 62]) and track length to further investigate how sampling time, spatial resolution, optic flow, and global information content in the environment interact to direct attention and memory during goal-directed behavior.

More broadly, we suggest that the presence or absence of a response to a particular stimulus could be interpreted as whether that dimension of experience was incorporated in memory. It is known that attention, task engagement, and learning exert profound influences on the decodability of sensory and spatial variables from hippocampal activity [4, 5, 28, 30, 47, 63]. The mechanisms that underlie this selective filtering of experience remain poorly understood and likely involve myriad upstream computations that occur between the primary sensory cortices and the hippocampal circuit [7, 64]. The advent of high density, multiregional neural recordings [7, 65, 66] will enable future work to refine the precise transformations that drive the confluence of sensory and action related coding in the hippocampus [62], and how and where contextual modulation of information transmission is implemented. Notably the task-relevance of stimuli has been shown to modulate neural responses in the cortex [67–70], which could contribute to filtering certain information streams during the formation of a memory trace in the hippocampus. Recordings of entorhinal inputs to the hippocampus indicate that there are pervasive sensory and self-motion signals present in the direct inputs to CA1 pyramidal neurons [57, 71, 72]. This convergence may be exploited by local circuit mechanisms in the hippocampus that serve to amplify certain environmental features or salient events [28, 72–78], particularly in response to novelty [42], which may also involve extrinsic and neuromodulatory control of excitability, plasticity, and neural assembly membership [72, 79–81].

Task engagement [30] and the task-relevance of stimuli impact hippocampal coding [4, 5, 27, 47, 63], but other salient stimuli and events can effect hippocampal neural activity despite no clear relationship to behavioral goals or reward [6–9, 78]. Considering that unsupervised learning is widespread in the dorsal cortex [82], we suspect that hippocampal codes remain highly plastic during unstructured experience. Even during performance of cognitive tasks where the parameters are known to the experimenter, it is always possible that the animal holds a different understanding or other uncontrolled motivations that impact what dimensions of experience are remembered.

## Acknowledgments

We thank C. Petersen for critical contributions to equipment; C. Petersen, W. Gerstner, and members of the Petersen and Gerstner labs for insightful discussions; S. Gex for hardware fabrication; and J. Bowler, J. Brea, and Z. Liao for comments on an earlier version of the manuscript. J.B.P. is supported by the EPFL ELISIR scholarship, the Fondation Marina Cuennet-Mauvernay, and a Novartis Foundation Young Investigator Grant. D.M. is supported by a Boehringer Ingelheim Fonds PhD fellowship.

## Author Contributions

O.C.U. and J.B.P. conceived the project and designed the experiments. All authors developed the VR system. O.C.U. collected and analyzed the data, supervised by J.B.P. O.C.U. and J.B.P. wrote the paper with input from all authors.

## Declaration of Interests

The authors declare no competing interests.

## Methods

### Resource Availability

Further information and requests for resources and reagents should be directed to the corresponding author James B. Priestley.

### Materials Availability

This study did not generate new unique reagents.

### Data and Code Availability

- Imaging data will be deposited on the DANDI archive and made publicly available as of the date of publication.
- All original code will be deposited at Zenodo and made publicly available as of the date of publication.
- Any additional information is available from the lead contact upon reasonable request.

## Experimental Model and Subject Details

All procedures were approved by Swiss Federal Veterinary Office (License number VD3984) and were conducted in accordance with the Swiss guidelines for the use of research animals. Experiments were performed with 7 adult (8-16 weeks) male and female C57Bl/6 mice (Charles River Laboratories, France).

## Method Details

### Behavior and Imaging

#### Viruses

Pyramidal cell imaging experiments were performed by injecting a recombinant adeno-associated virus (rAAV) encoding *GCaMP8m* (rAAV9-*Syn-GCaMP8m-WPRE-SV40*, Addgene) into male and female wild-type mice.

#### Surgical procedure

Methods for viral delivery and surgical implant of imaging window and headposts were largely identical to previous work [42]. Briefly, mice were anesthetized under isofluorane and the virus was injected in dorsal CA1 (-2.1 mm AP; -1.6 ML, -1.2 DV relative to bregma; 500 nL) using a Nanoject syringe. Mice recovered in their home cage for 3 days following viral infusions. We then gently aspirated the cortex overlying the left dorsal hippocampus and implanted a 3 mm glass-bottomed stainless steel cannula for imaging access, and cemented a titanium headpost to the skull for head-fixation. For all surgeries, monitoring and analgesia (ibuprofen) was continued for 3 days postoperatively.

#### Behavioral apparatus and virtual reality system

Mice were head-fixed above a low-friction, light weight running wheel [42, 43, 83]. The axle of the running wheel was coupled to a rotary encoder (Yumo COM-11102), which was connected to an Ardunio UNO that decoded and buffered the quadrature data from the encoder and transmitted these position updates to the VR control computer on each update cycle of the VR (≈ 60 Hz). To deliver water rewards to the animal, a stainless steel feeding needle (Fine Science Tools) was fixed on a custom 3D printed holder in front of the wheel and connected by silicon tubing to a gravity-fed water dispensing system gated by a solenoid valve (LHDA0531115H, the Lee Company). Licks were detected by soldering an insulated wire to the feeding needle, which was connected to an Adafruit CAP1188 capacitive touch sensor board. We used a second Arduino UNO connected to the touch sensor and solenoid valve by a custom PCB to monitor lick contacts and control the reward dispensing system. Experiments were managed through custom software built with the Unity video game engine [42, 43, 84], which interfaced with the two Arduino UNOs to monitor and actuate the physical elements of the VR behavioral tasks (real-time updating of virtual position with physical movement, detection of licks, and reward delivery) and synchronization with the 2-photon imaging system (ScanImage, MBF Biosciences). All system elements were connected by USB and communicated via custom serial protocols.

The running wheel was surrounded by 5 24” IPS display panels (U2424H, Dell) arranged in a half-octagon, covering approximately 225° of the visual field of the mouse. The monitors were connected to the VR control computer, which rendered the VR scene across the 5 displays, and displayed behavioral monitoring information on a 6th monitor outside of the VR rig. The entire VR screen array was mounted on a vertically movable frame that was counterbalanced on a gantry surrounding the microscope, so that it could be easily raised to access the animal and the microscope.

#### Procedural generation of virtual reality environments

The visual textures applied to the walls of the virtual tunnels were generated using the improved Perlin noise algorithm [40, 41], a procedural noise process that produces approximately statistically uniform, locally smooth random patterns by interpolating between randomly sampled displacement vectors on a uniformly spaced lattice. The spacing of the lattice grid determines the dominant frequency characteristic and spatial autocorrelation of the texture, allowing us to create environments of Low or High visual complexity (Fig. 1). The noise algorithm was implemented directly in Unity as a shader effect, from which 2048 × 256 pixel PNG textures were sampled (Fig. 1B). To increase the contrast of the resulting textures, eight instances (“Octaves”) of the noise process were multiplied while keeping the relative frequency (“Lacunarity”) and amplitude (“Persistence”) constant (fixed at a value of 1).

#### Behavioral training

Starting 7 days after the implant surgery, mice were habituated to handling and head-fixation as previously described [42]. After two days of acclimation and free running on the wheel, we began exposing the animal to a 4 m virtual environment for training. The training environment was also procedurally generated as described above, using visual textures of an intermediate frequency between the High and Low textures studied in the main experiments. When the mice reached the end of the virtual track, the screens were momentarily blanked for a 2 sec inter-trial interval (ITI), after which they were instantly teleported back to the beginning of the track at the start of the next trial. The animals’ line-of-sight was limited in all virtual environments by in-game lighting effects, so that only visual cues up to 1 m ahead of the animal were visible at each position.

At this point, mice were water deprived to 85-90% of their starting weight. Over a period of 1-2 weeks, we trained mice to run forward through the training environment and lick for small volume sucrose solution rewards (5% sucrose, ∼ 5µm per reward). Rewards were initially dispensed at random locations in the latter half of the environment on each trial (i.e. to habituate the animal to the range of distances used for the reward locations in the eventual context discrimination experiments) to encourage running. Rewards were initially automatically dispensed whenever the animal crossed the boundary of the reward zone on each trial. During training, we gradually reduced the probability of automatic rewards to zero while introducing lick-dependent rewards (i.e. the reward is only dispensed if the touch sensor detects a lick within the boundaries of the reward zone), as mice developed preparatory licking behavior. Mice reached training criteria when they could consistently run > 100 trials in under an hour and collect sufficient water rewards to maintain > 85% body weight. Mice were given additional water as needed daily to maintain weights.

#### Context discrimination paradigm

For context-discrimination experiments, mice ran through pseudo-randomly alternating trials (maximum of 2 repeats of the same environment) in two procedurally generated environments (see above), each populated with either High or Low frequency Perlin noise patterns (Fig. 1). The number of trials in each environment was approximately balanced across the session. For each mouse, the goal zone was randomly assigned in the High and Low contexts to either a Near (2.5 m) or Far (3.4 m) distance from the start of the trial. This mapping between High/Low and Near/Far was maintained for each mouse in the subsequent perturbation experiments (Replaced and Inserted trials, Fig. 4, 5). In Replaced trials, the absolute distances to Near and Far goals from the beginning of the trial were identical to the distances in Original trials, although the sensory cues at the goal locations were changed. In Inserted trials, the distances to goals was effectively increased by 2 m in both cases (Near = 4.5 m, Far = 5.4 m), but was colocalized with same sensory cues present at the goals in Original trials. At the beginning of the experiments, both the environments and the discrimination task were novel to the animal. Mice typically learned the Original two-context discrimination task within a single session. To avoid contamination by transient learning-related effects, we analyzed neural data during expert sessions in each mouse (namely, after 1 or 2 prior sessions in the discrimination task).

#### Procedural environment perturbations

To construct the perturbed virtual environments (Fig. 4, 5, 6), a new noise texture was sampled using the same parameters but varying the random seed used to draw the displacement vectors. For the environments in Replaced trials (Fig. 4), a new texture was sampled and the second half of the Original texture was replaced by the newly generated texture. For Inserted trials (Fig. 5, 6), a separate new texture was sampled as above and spliced into the middle of the Original texture (effectively extending the texture to a size of 3072 × 256 pixel). The second half of the Original texture became the final 3rd of the new, spliced texture, with the new texture inserted at the inner 3rd. In all cases, the manipulated sensory cues were not visible to the animal at the beginning of the trial (i.e., the track appears identical to Original trials) due to the limited line-of-sight described above. Original and perturbed trials could only be disambiguated once the animal had traveled farther into the environment.

#### 2-photon microscopy

Mice were habituated to the imaging apparatus (e.g. microscope/objective, laser, sounds of resonant scanner and shutters) during the training period. All imaging was conducted using a custom 2-photon microscope (Independent NeuroScience Services) equipped with an 8 kHz resonant scanner (Cambridge Technology), and a 16x NIR water immersion objective (Nikon, 0.8 NA, 3 mm working distance) or 10x air objective (Thorlabs, 0.5 NA, 7.1mm working distance). For excitation, we used a 920 nm laser (50-100 mW at objective back aperture, Spectra-Physics). Green (GCaMP8m) fluorescence was collected through an emission cube filter set to a GaAsP photomultiplier tube detector (Thorlabs, PMT-2101). All experiments were performed at 1.0-2.5 digital zoom, acquired as 512 × 512 pixels images at 30 Hz.

## Quantification and Statistical Analysis

### Behavioral data analysis

Behavioral data (velocity, position, licks, reward dispense events, VR scene changes, image synchronization signal) were logged at each VR update cycle (≈ 60 Hz). For all joint analyses with imaging data, behavioral data was aligned and downsampled to synchronize with each imaging frame.

### Task engagement analysis

Mice tended to disengage from the behavioral task toward the end of the session [30], likely after reaching satiety. We fit a 2-state hidden Markov model to the sequence of goal zone velocity and licking data (proportion of trial licks that occur in the goal zone) on each trial (i.e. each observation was a 2-element vector encoding these behavior data for a single trial, modeled as independent Gaussians, and the sequence length was equal to the number of trials). We labeled the latent states post hoc as Engaged and Disengaged, based on the different observation statistics (i.e., the Engaged state corresponded to low velocity and high licking in the goal zones, Fig. S1). The state sequence vector for each session was calculated via the Viterbi algorithm [85] and a cutoff trial was established by locating the end of the last contiguous block of ≥ 10 engaged trials. We discarded the remaining trials in the session from further analysis. Using this procedure, we found that mice typically completed ≈ 200 trials before disengaging from the task, which corresponded to ≈ 1 mL of water rewards.

#### Velocity and licking response curves

Velocity was calculated from the rotary encoder position updates, converted to cm/sec, and smoothed in time with a Gaussian filter (*σ* = 83*ms*). The spatial profile of velocity on each trial was calculated by binning and averaging velocity at each position. This was then used to calculate the peri-goal velocity curves (Fig. 1, 4, 6). We similarly estimated the animals’ licking rate at each position.

#### Decoding trial type from behavior

We used logistic regression to predict trial identity (High vs Low context in Fig. 1, Original vs Replaced/Inserted in 4, 6) using the average velocity and lick rate in the 28 cm (7 spatial bins) before the leading edge of the goal/non-goal zone on each trial. For trial type decoding, we balanced the number of trials in each class by discarding trials from the beginning of the session for the more numerous trial type, and then computed decoding accuracy via 10-fold cross-validation using balanced training and test sets.

#### Comparing neural and behavioral separability

Neural and behavioral decoding accuracy was compared via Pearson correlation (Fig. 4K).

### Neural data analysis

#### Image preprocessing

Imaging data was motion corrected and ROIs detected using Suite2p (in “sparse mode”) [44], followed by Suite2p’s standard fluorescence extraction and neuropil correction.

Identified ROIs were curated post-hoc using Suite2p’s graphical interface to exclude non-somatic components.

#### Event detection

All fluorescence traces were deconvolved to detect putative spike events, using the OASIS implementation of the fast non-negative deconvolution algorithm [45]. As in Priestley et al. [42], Ahmed et al. [63], we discarded any events whose amplitude was below 4 median absolute deviations of the raw trace, and binarized the resulting signal for all subsequent analysis to indicate whether the neuron was active in a given frame. We note that we do not claim to uncover true underlying spike times, but rather use deconvolution as a tool for denoising and reducing the impact of the calcium autocorrelation on our analysis.

#### Calculating spatial tuning curves

Spatial tuning curves were calculating for each neuron on each trial. The virtual track was discretized into 4 cm bins (i.e. 100 bins in 4 m environments; 150 bins in 6 m environments), which were used to compute a histogram of neural events. After normalizing for animal occupancy, the histogram was convolved with a Gaussian kernel (*σ* = 8 cm / 2 bins) to obtain a smooth activity rate estimate.

#### Decoding context

We decoded context (the environment identity) by independently training a linear SVM classifier at each position (or goal-relative position) in the environment (Fig. 1, 3, 4. Decoding results are reported as the cross-validated accuracy, obtained by randomly splitting the trials into equal sized training and test sets. Cross-validated accuracy was computed as the average of 10 randomized training/test splits. The training/test splits were synchronized across the independent classifiers at each position, which allowed us to additionally compute aggregate trial-level prediction of the context by taking the majority vote of the context classifiers at each position, for each trial in the test set.

#### Spatial tuning score

We computed a pseudo-probability mass function *p*(*x_i_*) for each spatial tuning curve, where *x*_1_, *x*_2_, …*x_n_* are the discrete position bins, by normalizing the tuning curve so that it sums to 1 over positions. We then compute the un-normalized tuning score *s*^ as the KL-divergence between *p* and the uniform distribution *u*:

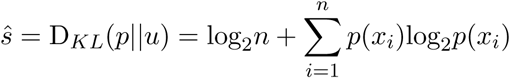

Intuitively, uniform activity over space will give the minimum score *s* = 0, while having all activity concentrated in a single position bin will yield the maximum score of *s* = log_2_(*n*). Inhomogenous spatial tuning can also arise simply from very sparse, noisy activity, a regime in which using raw information-theoretic metrics can give misleading results [86]. To control for the effects of firing rate and sampling, we used a normalized tuning score *s* by standardizing *s*^ relative to a null distribution:

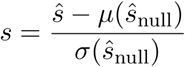

The null distribution was sampled by calculating the tuning score on a distribution of null tuning curves constructed by circularly shifting neural activity on each trial independently by a random position offset to disrupt trial-to-trial spatial correlations.

#### Decoding spatial position

In order to assess position coding and context discrimination, we trained a decoder to predict the spatial location of the animal from neuronal population activity, using an ensemble of support vector machines (SVM, “one-vs-one” multi-class decoding [42, 63]). Spatial results were similarly cross-validated across 10 repeats of randomized 50/50% splits of trials into training and test sets. We assessed cross-validated performance of each classifier as both the discrimination accuracy of the underlying one-vs-one classifiers at every possible distance in the environment (Fig. 2E, F) and the mean absolute error of the ensemble multi-class predicted positions (Fig. 2G).

#### Decoding goal-relative position and context

In Fig. 3, we repeated our spatial and context decoding analyses in goal-aligned coordinates. Namely, we cropped and aligned the spatial tuning curves of neurons in both contexts so that the analysis included an equivalent amount of distance before and after the goal zone in each context (2 m before and 0.56 m after). Cross-validation was performed as described above.

#### Cross-condition generalization performance

We assess the generalization of goal-relative distance coding across contexts [11, 12, 27] using cross-condition generalization performance (CCGP, [46]): we trained classifiers to predict position (relative to goal) in one context (High/Low, or equivalently, Near/Far) and then tested the performance in the opposite context (Fig. 3E,G). If the representations in each context converge to a common goal/action code or are arranged in parallel in the neural state space to abstractly encode the goal distance (Fig. 3A), we would expect high CCGP (either specifically around the goal location for reward cells [11], or throughout the environment [12, 27].

#### Population vector spatial cross-correlations

We compared population vector similarity across pairs of positions, both within environment and across environments, using Pearson’s correlation (Fig. 3, 4, 5, 6). For comparisons within the same environment, these correlations were calculated as the population vector correlation between pairs of positions in even vs odd trials. The Δ correlation matrices in Fig. 4 and 5 were formed by calculating the difference between the the even/odd correlations in Original trials and the Original/Replaced (Inserted) trials. In Inserted trials, this latter cross-correlation matrix is not square (comparing a 4 m vs 6 m environment). We calculated the Δ matrix first by subtracting the even/odd Original matrix from the first 4 m square of the Original vs Inserted matrix. We the calculated the Δ for the last 2 m by subtracting the rightmost column (last 2 m) of the Original matrix from the rightmost column (last 2 m) of the Original vs Inserted matrix.

## Supplementary Materials

**Figure S1.**
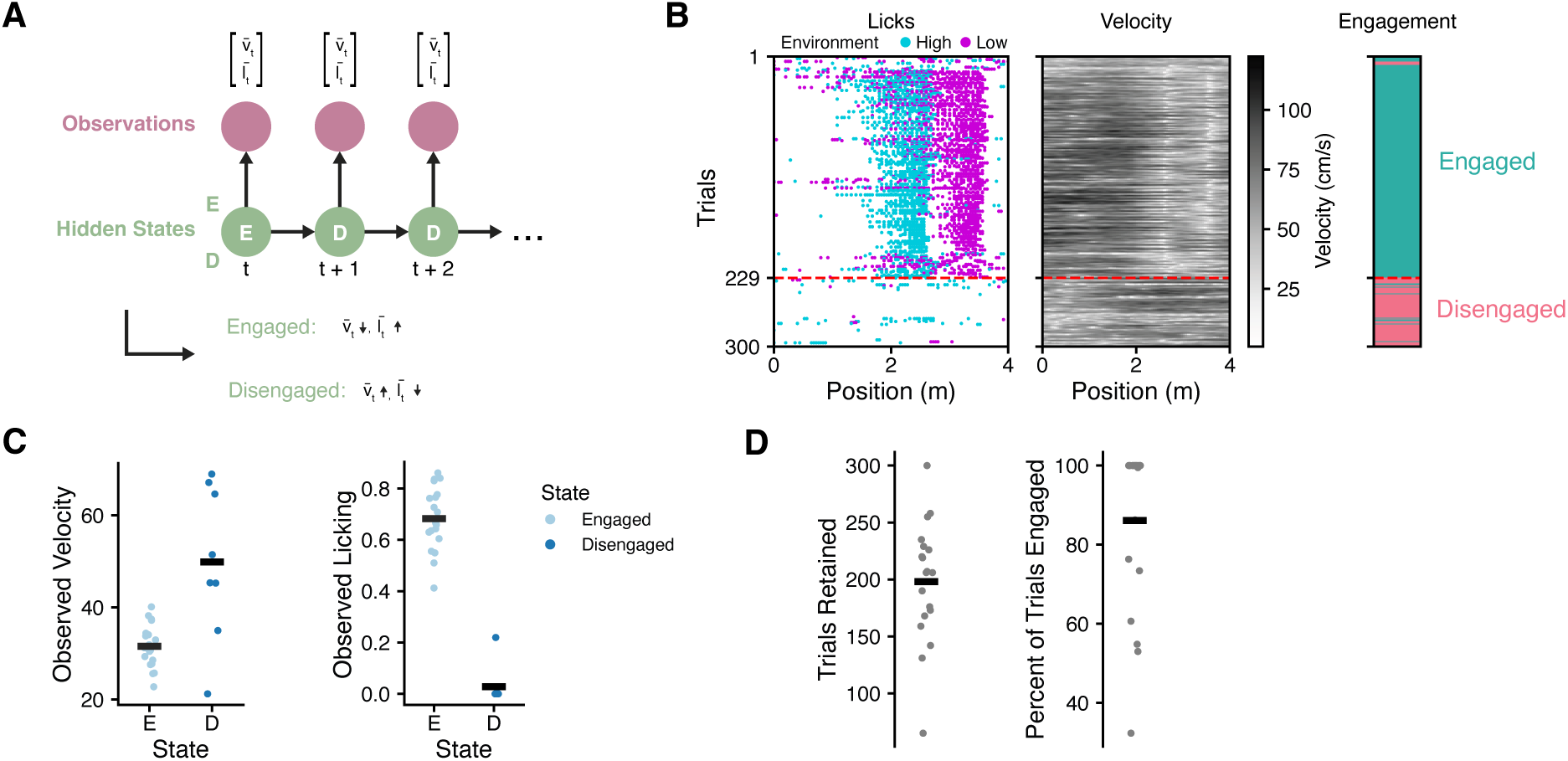
Classifying changes in task engagement. **A:** Schematic of hidden Markov model (HMM) used for task engagement classification. The observation variables consisted of the mean velocity *v̅t* in the reward zone and the proportion of licks within the reward zone *l̅t* in each trial. The model switched between two latent states, which we post hoc identified as Engaged (E) and Disengaged (D) based on the learned observations. **B:** Example raster plot of licks (left), trial-by-trial velocity (middle), and latent states predicted by the HMM (right) for an example session. The dotted red line indicates the cutoff trial for the session, defined as the end of the last contiguous block of *≥* 10 trials assigned the Engaged state by the HMM. **C:** Number of trials retained after discarding trials after the disengagement threshold in each session. **D:** Percent of trials classified as Engaged for each session.

**Figure S2.**
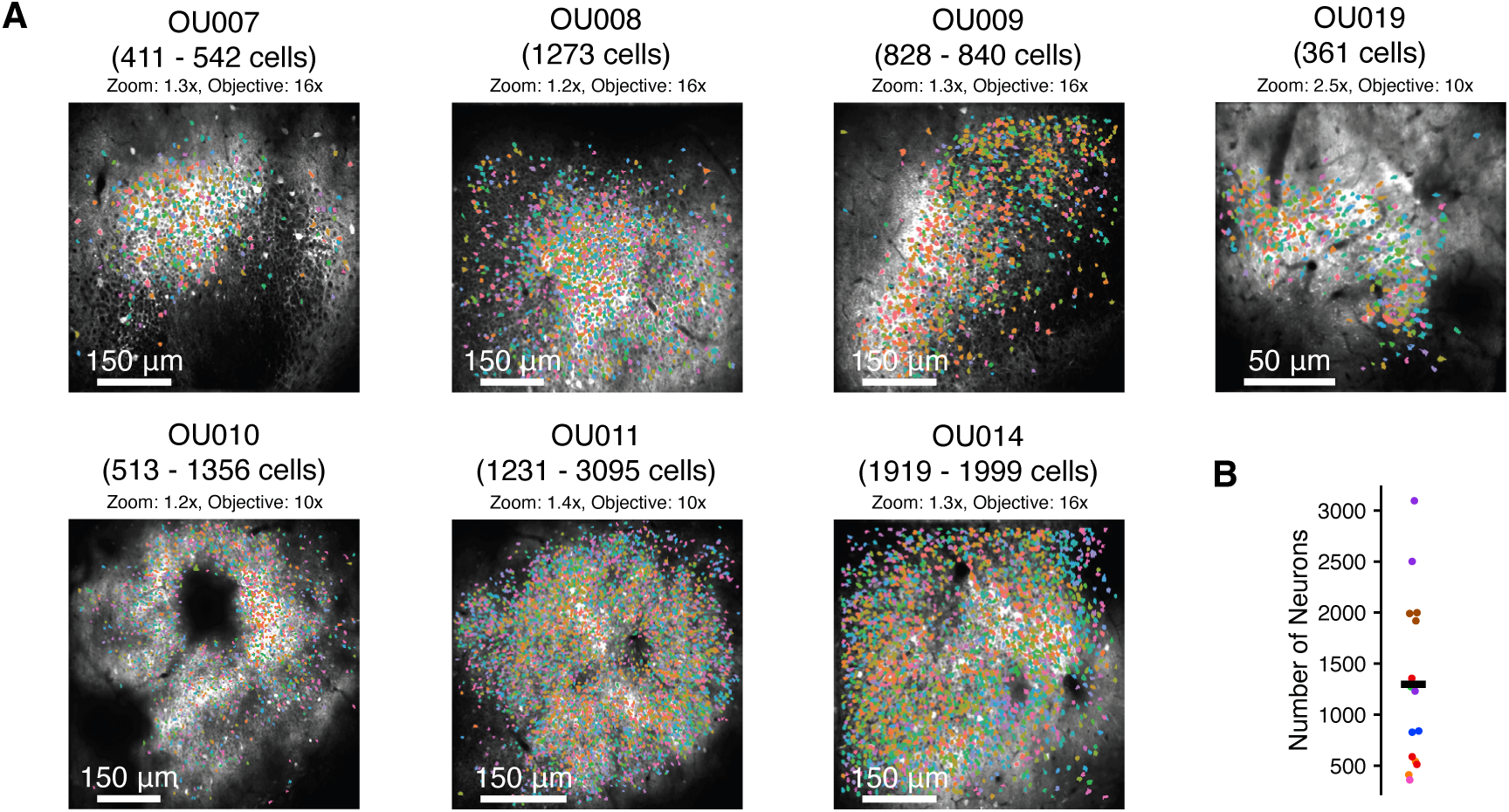
Imaging data overview. **A:** Example field-of-view (FOV) from each mouse in the dataset, with segmented ROIs overlaid. **B:** Number of segmented neurons per session. Colors correspond to individual mice.

**Figure S3.**
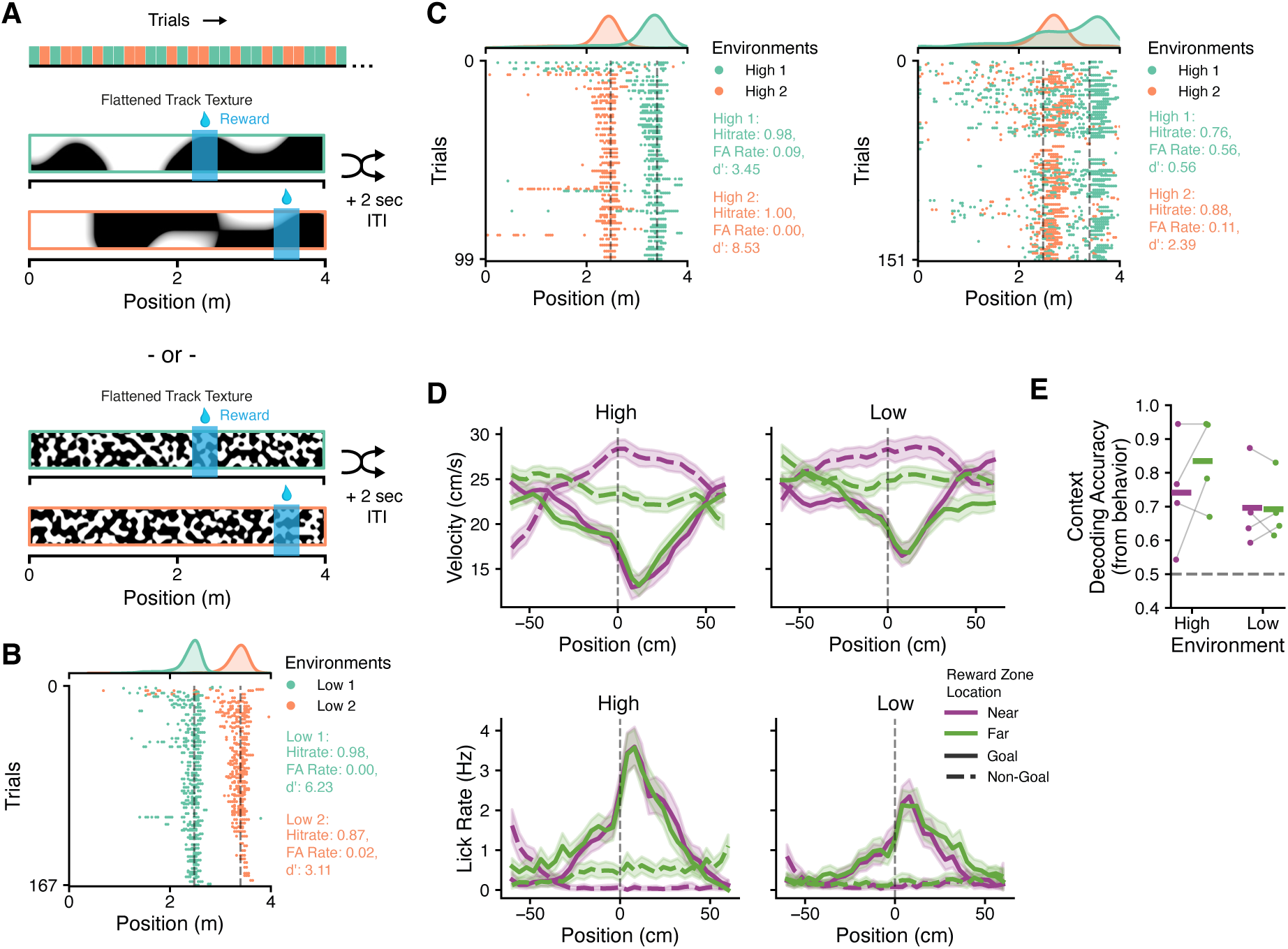
Mice can discriminate two procedural VR environments of equivalent visual complexity. **A:** Schematic of behavioral task. Mice learned to discriminate between two VR environments that were generated with the same procedural noise parameters (Low vs Low, middle, or High vs High, bottom). As in the main experiments, context discrimination was measured by the restriction of licking to one of two possible goal zones placed at different distances in each environment. Trials pseudorandomy alternated between the two contexts as before (top). **B:** Example trial-by-trial licking data for an example Low vs Low discrimination session. The mouse reliably discriminates between the two environments. **C:** As in **B**, but for two example High vs High sessions, one with highly accurate discrimination (left) and one with poor discrimination (right). **D:** Velocity (top) and licking (bottom) behavior around the goal (non-goal) zones in High vs High (left) and Low vs Low (right) discrimination sessions. Colors indicate the goal location (i.e. context identity). **E:** Logistic regression to predict trial type (Context 1 or Context 2) from behavior (pre-goal licking and velocity), shown separately for behavior around the Near (purple) or Far (green) distance, in High vs High and Low vs Low sessions. While performance is variable, some mice reliably differentiate the two High environments.

**Figure S4.**
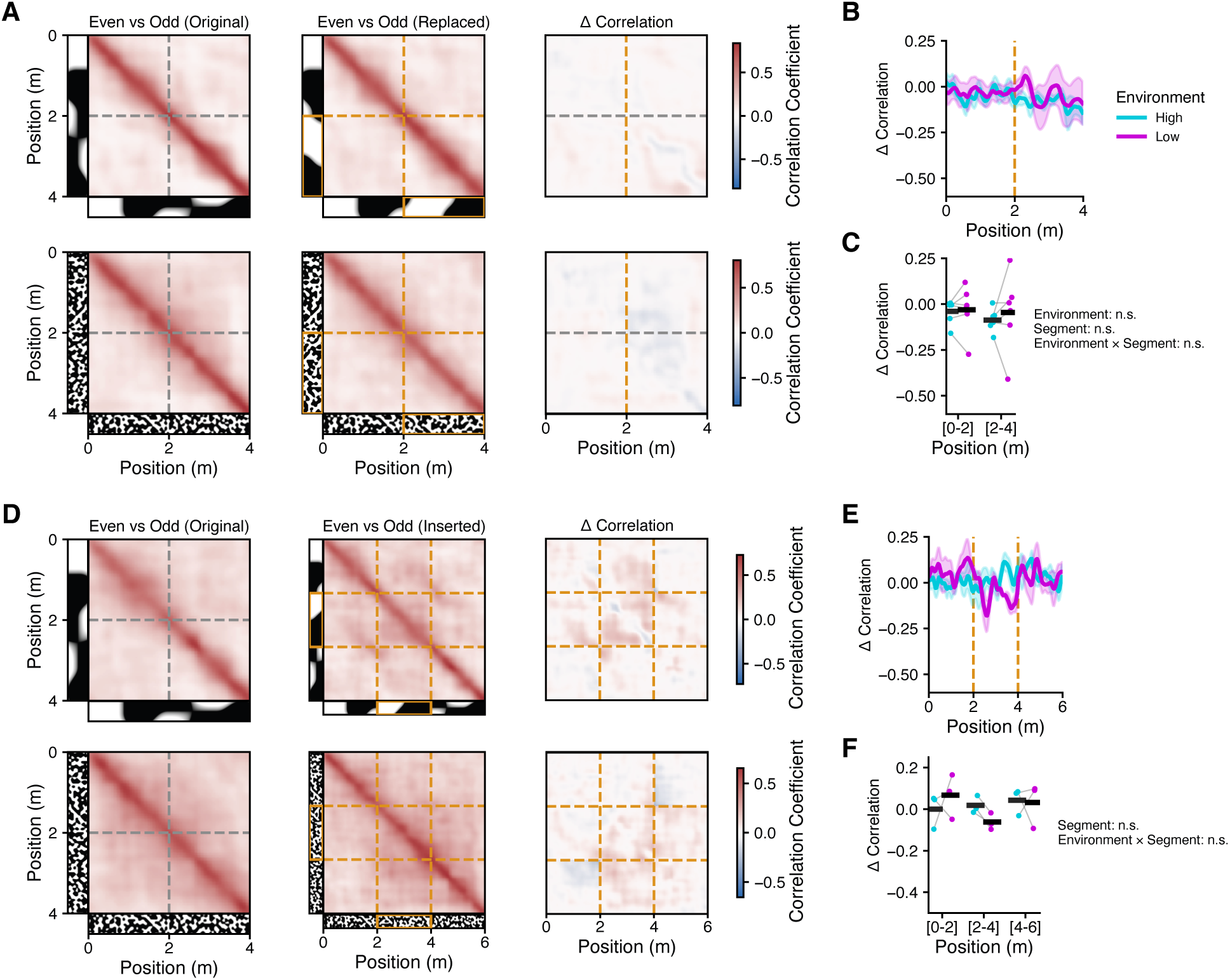
Perturbed environments produce similar quality place codes. **A:** Population vector cross-correlations for each pair of spatial positions, between Even vs Odd trials in the Original environment (left), between Even vs Odd trials in the Replaced environment (center), and their difference (right), shown for the Low (top) and High (bottom) contexts. Both Original and Replaced environments form reliable sequences. **B:** Summary of the diagonal of the Δ correlation matrices plotted in **A** (right) for each mouse. **C:** Average Δ correlation in **D** for each mouse and context, shown for the first half (0-2 m) and second half (2-4 m) of the environment (linear mixed effects model). **D-F**: As in **A-C**, for Inserted experiments. Δ correlations for [4, 6] segments in Inserted were computed compared to [2, 4] in Original trials.

## Notes

### Competing Interest Statement

The authors have declared no competing interest.

